# MEK1/2 and ERK1/2 mediated lung endothelial injury and altered hemostasis promote diffuse alveolar hemorrhage in murine lupus

**DOI:** 10.1101/2024.05.07.593006

**Authors:** Haoyang Zhuang, Shuhong Han, Neil S. Harris, Westley H. Reeves

## Abstract

**Objective:** About 3% of lupus patients develop severe diffuse alveolar hemorrhage (DAH) with pulmonary vasculitis. B6 mice with pristane-induced lupus also develop DAH, but BALB/c mice are resistant. DAH is independent of TLR signaling and other inflammatory pathways. This study examined the role of the mitogen-activated protein kinase pathway (MEK1/2-ERK1/2, JNK, p38).

**Methods:** B6 and BALB/c mice were treated with pristane ± inhibitors of MEK1/2 (trametinib/GSK1120212, “GSK”), ERK1/2 (SCH772984, “SCH”), JNK, or p38. Effects on lung hemorrhage and hemostasis were determined.

**Results:** GSK and SCH abolished DAH, whereas JNK and p38 inhibitors were ineffective. Apoptotic cells were present in lung from pristane-treated mice, but not mice receiving pristane+GSK and endothelial dysfunction was normalized. Expression of the ERK1/2-regulated transcription factor *Egr1* increased in pristane-treated B6, but not BALB/c, mice and was normalized by GSK. Pristane also increased expression of the anticoagulant genes *Tfpi* (tissue factor pathway inhibitor) and *Thbd* (thrombomodulin) in B6 mice. The ratio of tissue factor (*F3*) to *Tfpi* increased in B6 (but not BALB/c) mice and was normalized by GSK. Circulating Thbd protein increased in B6 mice and returned to normal after GSK treatment. Consistent with augmented endothelial anticoagulant activity, pristane treatment increased tail bleeding in B6 mice.

**Conclusion:** Pristane treatment promotes lung endothelial injury and DAH in B6 mice by activating the MEK1/2-ERK1/2 pathway and impairing hemostasis. The hereditary factors determining susceptibility to lung injury and bleeding in pristane-induced lupus are relevant to the pathophysiology of life-threatening DAH in SLE and may help to optimize therapy.

## Introduction

Severe diffuse alveolar hemorrhage (DAH) is one of the clinical manifestations of pristane-induced lupus (1–3). C57BL/6 (B6) mice are susceptible to DAH while BALB/c and most other strains are resistant (1). Hemorrhage is accompanied by endothelial injury, ER stress, and an ANCA-negative pulmonary capillaritis closely resembling the pathology of DAH in systemic lupus erythematosus (SLE) patients (1–4). Although only 3-4% of SLE patients have life-threatening hemorrhage, focal lung hemorrhage is seen at autopsy in 30-66% of SLE patients, suggesting a high prevalence of subclinical disease (5). DAH also develops in antiphospholipid syndrome, rheumatoid arthritis, and antineutrophil cytoplasmic antibody (ANCA)-associated vasculitis (6) and it can complicate therapy with rapamycin (sirolimus) (7). Rapamycin treatment exacerbates DAH in pristane-treated B6 mice (S Han, et al. submitted). Pristane-induced DAH is mediated by monocytes and macrophages (ML), complement component C3, and CR3 (CD11b/CD18), but does not require neutrophils (2). Mice lacking Tlr7, MyD88, TRIF, the type I interferon receptor (IFNAR), or TNFα are fully susceptible (2, 8, 9). Thus, DAH is a TLR, IFNAR, and TNFα -independent clinical manifestation of lupus with a striking strain-specificity. Understanding the pathogenesis of DAH in B6 vs. BALB/c mice may shed light on why only 3-4% of SLE patients develop life-threatening DAH, whereas 30-66% have mild disease, and the rest are resistant.

The inflammatory pathway(s) mediating DAH in pristane-induced lupus are unclear. Although DAH is independent of TLR, inflammasome (caspase-1), and Nos2 signaling (2), a variety of other germline-encoded pattern recognition receptors converge on the mitogen-activated protein kinase (MAPK) and nuclear factor kappa B (NFκB) pathways, promoting inflammation (10). As many as 14 MAPKs have been reported in mammalian cells, most notably extracellular signal-regulated protein kinase 1 (ERK1), ERK2, p38, Jun N-terminal kinase 1 (JNK1), and JNK2 and their upstream activators, which link MAPK activation to innate immune receptors (10).

The objective of the present study was to define the role, if any, of MAPK activation in pristane-induced DAH. We found that inhibitors of ERK1/2 and its upstream activators mitogen-activated protein kinase kinase 1 (MEK1) and MEK2 block the induction of DAH and lung injury in B6 mice following pristane exposure. In contrast, inhibition of JNK or p38 was not protective. Susceptibility to DAH also was dependent on inter-strain disparities in hemostasis between B6 and BALB/c mice, which may help explain why only a small subset of SLE patients develops life-threatening DAH.

## Materials and Methods

### Mice

Mice were maintained under specific pathogen-free conditions. Female 8-12LweekLold C57BL/B6 (B6) and BALB/cJ mice were from The Jackson Laboratory. The animals were randomly divided into treatment or control groups. To induce lupus, 0.5 ml of pristane (SigmaLAldrich) was administered i.p. Controls were left untreated. Experiments were repeated at least twice. Peritoneal exudate cells (PEC) were collected 14-d after pristane treatment unless otherwise noted. DAH was assessed as described (2). After euthanizing the mice, bronchoalveolar lavage (BAL) was performed with 1 mL of phosphate-buffered saline.

Cells were pelleted and the undiluted supernatant was tested by ELISA (BioLegend) for the monocyte-attractive, Egr1-regulated, chemokine Ccl2. After BAL, the lungs were fixed in 10% neutral buffered formalin and embedded in paraffin. This study followed the recommendations of the Animal Welfare Act and US Government Principles for the Utilization and Care of Vertebrate Animals and was approved by the University of Florida Institutional Animal Care and Use Committee.

### Inhibitor treatment

B6 and BALB/c mice, age 8-12 weeks, were treated with pristane with or without the ERK1/2 inhibitor SCH772984 (Selleckchem, Houston TX, 15 mg/kg i.p. daily) or the MEK1/2 inhibitor GSK1120212 (trametinib, ApexBio, Houston, TX, 0.1 mg/kg orally every 2 days). SCH772984 (SCH) is a potent and highly selective ERK1 and ERK 2 inhibitor (11, 12). GSK1120212/trametinib (GSK) is a highly selective allosteric inhibitor of MEK1 and MEK2 that causes sustained inhibition of downstream MEK1/2-mediated ERK1/2 phosphorylation without significantly inhibiting 98 other kinases (13, 14). A group of mice was treated with AZD8330 (Selleckchem, 0.5 mg/kg/d oral gavage), a newly developed MEK1/2 inhibitor (15) and another group with ulixertinib (Selleckchem, 50mg/kg/d oral gavage), a potent, specific, and reversible ERK1/2 inhibitor (16). Additional mice were treated with pristane with inhibitors of p38 MAPK (ralimetinib/LY2228820 dimesylate, Selleckchem, 10 mg/kg/d oral gavage) or JNK1/JNK2/JNK3 (SP600125, Selleckchem, 15 mg/kg/d oral gavage), or with the cAMP analog dibutyryl-cAMP/bucladesine (APExBIO, 15 mg/kg/d i.p.). DAH was assessed at 14-d.

Ralimetinib/LY2228820 is a selective p38α and p38β inhibitor that potently blocks p38 MAPK signaling (17). SP600125 is a broad-spectrum inhibitor with 10-100-fold greater selectivity for JNK1, JNK2, and JNK3 vs. a variety of other kinases, including MKK4 (10-fold), ERK2 and p38 (100-fold), and certain serine/threonine kinases such as FLT3 (18). Dibutyryl-cAMP is a cell-permeable cAMP analog that preferentially activates protein kinase A (PKA), mimicking the action of endogenous cAMP (19). It also inhibits phosphodiesterase.

### Quantitative real time PCR (qPCR)

RNA was isolated from 10^6^ mouse PEC or lung tissue using TRIzol (Invitrogen). cDNA was synthesized using the Superscript II First-Strand Synthesis Kit (Invitrogen). SYBR Green qPCR was performed using a CFX96 thermocycler (Bio-Rad). Gene expression was normalized to 18S RNA, and relative expression was calculated using the 2^-ΔΔCt^ method. Primer sequences are listed in **Table 1**.

**Table 1.**
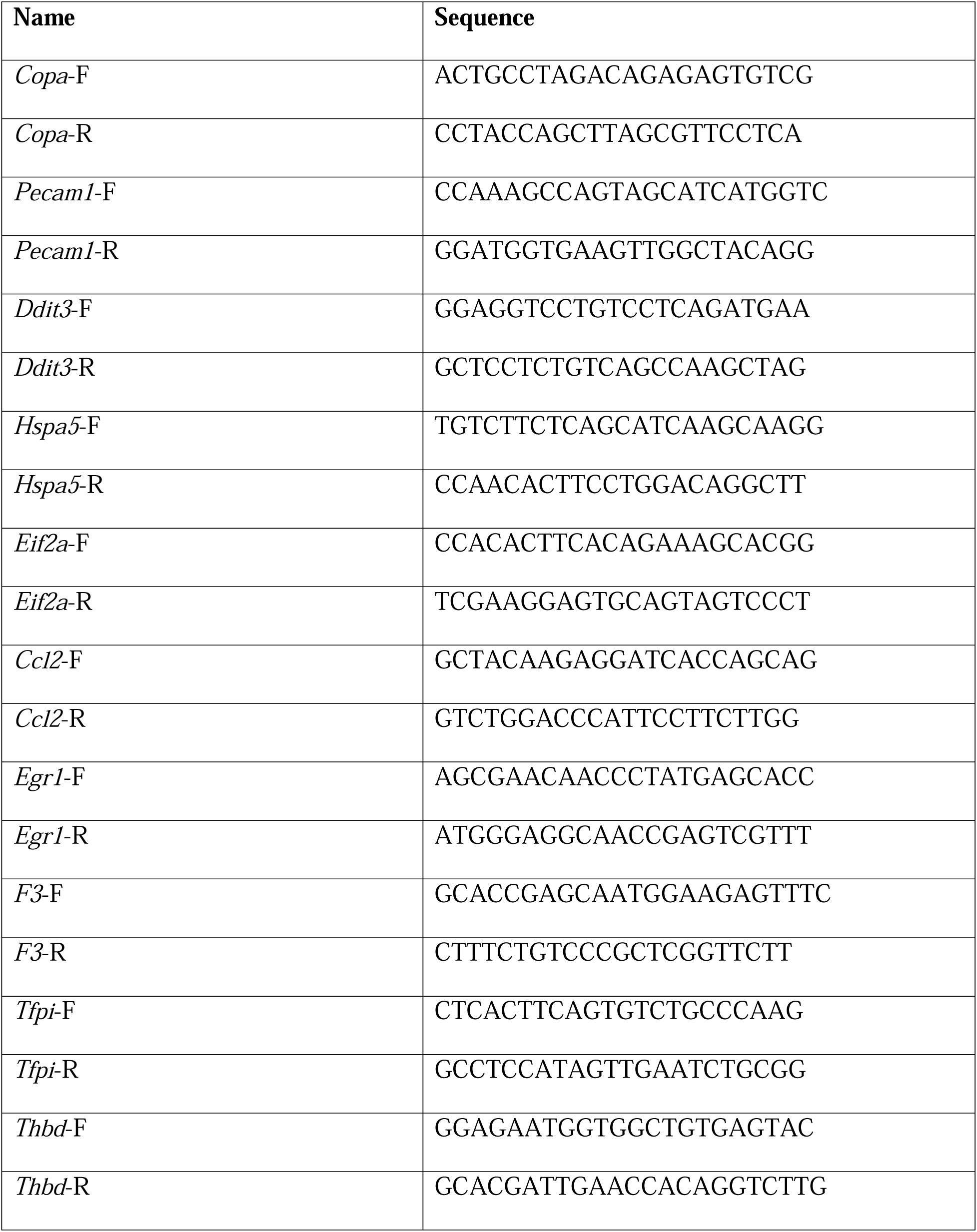
Primer sequences.

### Immunohistochemistry of lung tissue

Paraffin sections (4 µm) of formalin-fixed lung tissue obtained 14-d after pristane treatment were stained with H&E or Prussian blue. Single and double immunohistochemistry (IHC) were performed as described (20). Paraffin sections were dried on slides for 2-h at 60°C. Slides were placed in a Ventana Medical Systems immunostainer and deparaffinized. HeatLinduced epitope retrieval was performed with Ventana Medical Systems CC1 retrieval solution (30-min at 95–100°C). Primary antibodies anti-cleaved caspase-3 (1:100 Cell Signaling Technology, Danvers, MA), anti-CD31/Pecam1 (R&D Systems 1:200, Dako), and F4/80 (1:200, Fisher Scientific) were applied for 30-min at 37°C. For single IHC, primary antibodies were followed by peroxidaseLconjugated goat antiLmouse or goat antiLrabbit secondary antibodies (30 min). For double immunohistochemistry, incubation with peroxidase-conjugated anti-CD31/Pecam1 or anti-F4/80 was followed by incubation with alkaline phosphatase-conjugated anti-activated caspase 3 antibody. Reaction product was visualized using an ultraView DAB detection kit (brown) or an ultraView Universal Alkaline Phosphatase Red Detection Kit (red) (Vertana). Slides were counterstained with hematoxylin (Ventana Medical Systems).

### Dissociation of lung tissue

Mouse lungs were dissected into single lobes and rinsed in ice-cold PBS pH 7.2, then mechanically dissociated in enzyme mix for 30 min at 37°C following the manufacturer’s protocol (Lung Dissociation Kit, gentleMACS Tissue Dissociator, Miltenyi Biotec). The single cell suspension was applied to a 70 µm filter with 2.5 mL Separation Buffer, washed at 300×g for 10 minutes, and resuspended in flow cytometry buffer for flow cytometry.

### Flow cytometry

Flow cytometry was performed using anti-mouse CD16/32 (Fc Block, clone 2.4G2, BD Biosciences) before staining with primary antibody or isotype controls. Dissociated lung cells or peripheral blood (100 µL) was incubated 20-min in the dark with anti-CD45-allophycocyanin and anti-CD31-Pacific blue (Biolegend). Erythrocytes were lysed with Red Cell Lysis Buffer (Biolegend). After surface-staining, cells were fixed/permeabilized for 5-min (Fix-Perm Buffer, #00-5123-43, eBioscience), washed, and stained intracellularly for 20-min with anti-Bip-PE (Cell Signaling, Danvers, MA). The cells were washed and re-suspended in flow cytometry buffer and analyzed with a FACSymphony™ A5 flow cytometer. For annexin V staining, cells were incubated in 1:20 phycoerythrin-conjugated annexin V solution (BioLegend) for 15 min prior to flow cytometry following the manufacturer’s protocol.

### Hemostasis assays

Tail vein bleeding tests were performed on mice with or without pristane injection as described with minor modifications (21). Briefly, mice were anesthetized and 1.0-cm of the tip of the tail was removed with a sharp scalpel. The tail was immediately immersed into a 15-mL conical tube containing 10-mL 0.9% NaCl at 37°C and was removed after 60 sec. Heme content were measured using a Heme Assay Kit (Sigma-Aldrich) according to manufacturer’s protocol. Briefly, 50 µl of sample was incubated with 200 µl of Heme Reagent for 5 min at 22°C. Heme content was measured by the absorbance at 400 nm VERSA MAX microplate reader (Molecular Devices).

The prothrombin time (PT) and international normalized ratio (INR), which depend on factors VII, V, X, prothrombin, fibrinogen, and fibrin, and the activated partial thromboplastin time (aPTT), which depends on factors XII, XI, X, IX, VIII, V, prothrombin, and fibrin (22) were determined in B6 mice treated with pristane alone or pristane + GSK, or left untreated using an ACL TOP 750 coagulation analyzer (Werfen, Bedford, MA). A normal mouse PT is 9-12 s and normal aPTT is 20-30 s, but the values are strain-dependent (23).

### ELISA

Soluble thrombomodulin (sThbd) was measured at day 14 after pristane treatment in heparinized plasma by sandwich ELISA using a kit from Bioss (BSKM61670-96T) following the manufacturer’s protocol. D-dimer and ferritin were measured in serum using ELISA kits from MyBiosource (MBS269348 and MBS261944, respectively) following the manufacturer’s protocols.

### Statistical analysis

Data are representative of at least 2 independent experiments and are presented as mean ± SD. For normally distributed data, comparisons were performed using Student’s unpaired 2-tailed t-test (GraphPad Prism software, version 5). When data were not normally distributed, comparisons were analyzed by the Mann-Whitney U test. Frequency data were analyzed by Fisher’s exact test. Survival was analyzed by Kaplan-Meier test. One-way comparisons involving three or more groups were analyzed by ANOVA with Tukey’s or Dunnett’s multiple comparison test and Holm-Sidak correction or Kruskal-Wallis for non-normally distributed data with Dunn’s correction for multiple comparisons. P values less than 0.05 were considered significant.

## Results

Pristane-induced DAH and small vessel lung vasculitis are closely associated with microvascular injury and ER stress in the lung (3, 4) and can be prevented by monocyte/ML or complement depletion (2) or liver X receptor (LXR) agonist therapy (24). Conversely, DAH is exacerbated by enhancing ER stress (3). Identifying the inflammatory pathway that mediates lung injury has proved elusive. DAH is independent of TLR signaling, type I interferon, TNFα, inflammasomes, and nitic oxide signaling (2). Here, we examined the role of the three arms of the mitogen-activated protein kinase (MAPK) pathway (ERK, JNK, and p38). In particular, ERK1/2 and its upstream regulators MEK1 and MEK2 control early growth response 1 (Egr1), a transcription factor regulating coagulation, inflammation, vascular permeability, and apoptosis (25, 26). We treated mice with pristane alone or along with the highly selective MEK1/2 inhibitor trametinib (GSK1120212, “GSK”), which is FDA-approved for melanoma (27), or a selective ERK1/2 inhibitor SCH772984 (“SCH”). Therapy with either GSK (0.1 mg/kg p.o. daily) or SCH (25 mg/kg i.p. daily) arrested the onset of DAH and prevented mortality completely **(Fig. 1A)**. Initiation of GSK therapy as late as 6-d after pristane treatment was effective (not shown). In nearly all mice, there was no evidence of hemorrhage by gross pathology, H&E or Prussian blue staining for hemosiderin when the mice were treated with GSK or SCH along with pristane **(Fig. 1B-G)**. Moreover, although small blood vessels in the lungs from pristane-treated mice exhibited perivascular lymphocytic infiltrates consistent with vasculitis **(Fig. 1C)**, there was no evidence of pulmonary vasculitis in mice treated with pristane plus GSK **(Fig. 1D)** or SCH (not shown). A different MEK1/2 inhibitor (AZD8330, 0.5 mg/kg/d) and a different ERK1/2 inhibitor (ulixertinib) also blocked the onset of DAH and prevented mortality (not shown). In contrast, pristane-induced DAH was not prevented by treating the mice with drugs that inhibit p38 (ralimetinib/LY2228820) or JNK (SP600125) **(Fig. 1H)**. SP600125 is a broad-spectrum kinase inhibitor that blocks JNK activation 10-100 times more potently than other kinases (18). It inhibits ERK2 100-fold less potently than JNK, but had no effect on DAH. The anti-inflammatory cAMP analog dibutyryl cAMP/bucladesine also had little effect on DAH **(Fig. 1H)**. Thus, drugs that block the MEK1/2-ERK1/2 pathway (GSK, AZD8330, SCH) all prevented DAH, whereas drugs that inhibit p38 (ralimetinib/LY2228820) or JNK (SP600125) or activate cAMP (dibutyryl cAMP/bucladesine) did not.

**Figure. 1.**
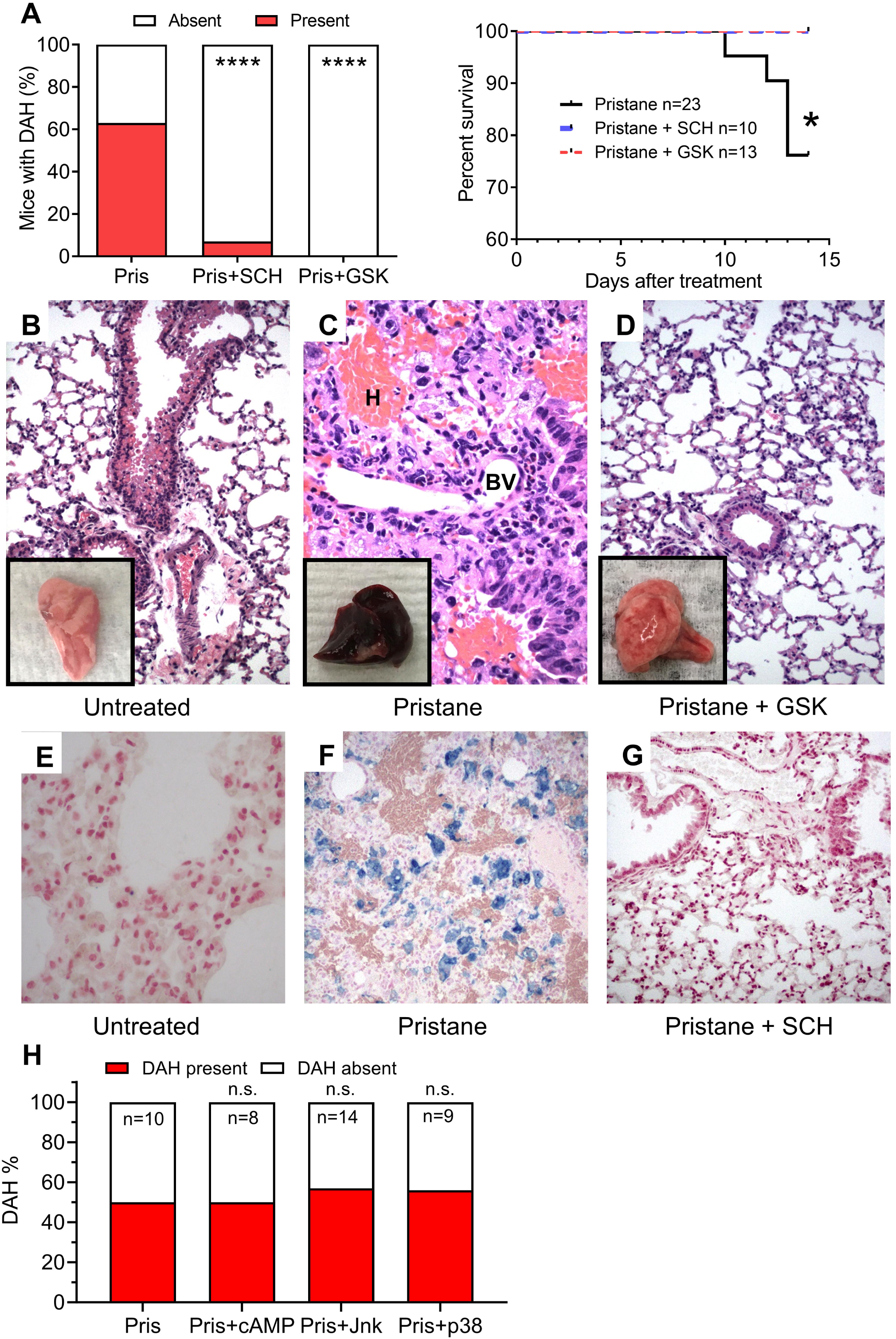
MEK1/2 and ERK1/2 inhibitors prevent DAH. **A)** B6 mice were injected i.p. with pristane (Pris) and 14-d later DAH was assessed. *Left*, Frequency of DAH in mice treated with Pris+SCH772984 (SCH, ERK1/2 inhibitor) or GSK1120212 (GSK, MEK1/2 inhibitor), or with pristane alone. *Right*, Survival of B6 mice treated with pristane, pristane + SCH, and pristane + GSK. * P < 0.05, **** P < 0.0001, t-test; n.s., not significant. **B-D)** Representative H&E staining and gross pathology (inset) of formalin-fixed lung from B6 mice treated for 14-d with pristane or pristane + GSK, or left untreated. BV, small blood vessel with vasculitis; H, hemorrhage. Original magnification X100. **E-G**, Representative Prussian blue staining (blue reaction product) for hemosiderin-laden macrophages in lung from B6 mice treated with pristane or pristane + SCH, or left untreated. Original magnification X100. **H,** B6 mice (n = 8-14 per group) were treated with pristane alone (Pris) or with pristane plus Jnk1/2/3 or p38 inhibitors. Or pristane plus a cAMP activator. DAH was assessed at d-14 after pristane treatment. n.s., not significant by Χ^2^.

GSK treatment also prevented apoptosis of pulmonary cells **(Fig. 2A-B)**. In comparison with control lung from untreated mice, lung tissue from B6 mice treated with pristane alone contained numerous cells with a positive immunohistochemical reaction for activated caspase-3 **(Fig. 2A)**. Activated caspase-3^+^ cells were not seen in lung from pristane-treated BALB/c mice or in lung from B6 mice treated with pristane + GSK. Analysis of dissociated lung cells by flow cytometry indicates that many lung endothelial cells from pristane-treated B6 mice are stained with anti-BiP (an ER stress marker) antibodies (3). However, double immunohistochemistry suggested that the activated caspase-3 positive cells in the lung tissue were not mainly CD31^+^ endothelial cells **(Fig. 2B)**. In lung tissue sections, most cells that stained positively for activated caspase-3^+^ were large with an irregular shape, but exhibited minimal staining with anti-F4/80, a ML marker **(Fig. 2B)**. Alveolar ML from mice treated with pristane + GSK were strongly F4/80^+^ but did not stain with anti-activated caspase-3 antibodies. Lung tissue also was stained with antibodies against CD31 (Pecam-1), a protein that maintains endothelial cell integrity (28, 29). The walls of blood vessels from untreated mice exhibited strong CD31 staining, whereas staining was greatly reduced in the walls of blood vessels from pristane-treated mice **(Fig. 2C)**. Treatment with GSK along with pristane restored the intense CD31 staining of small blood vessels in the lung.

**Figure 2.**
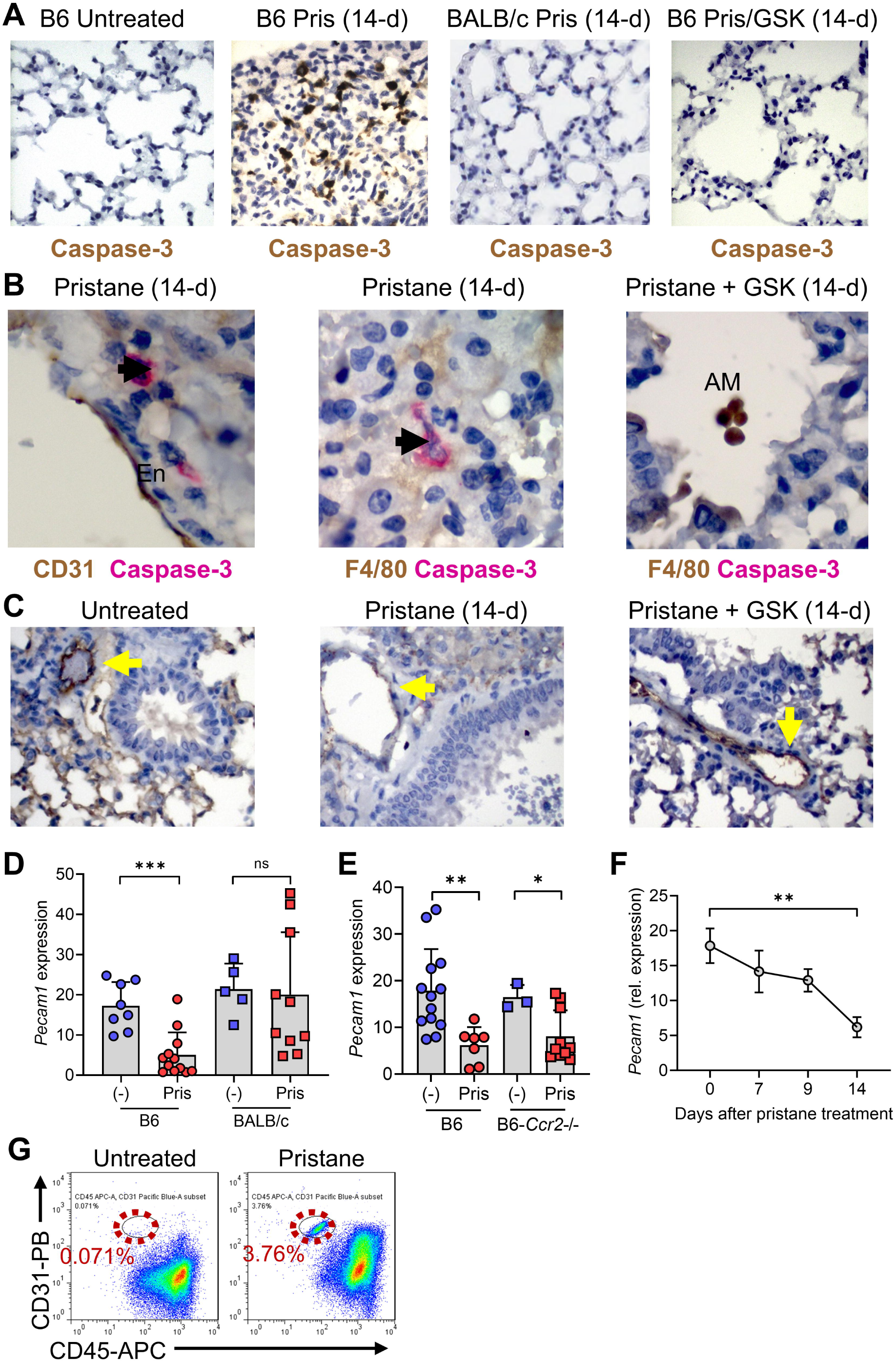
MEK1/2 inhibition prevents lung endothelial injury and apoptosis. **A,** Representative images of formalin-fixed lung tissue from untreated mice, or mice treated for 14-d with pristane alone or pristane+GSK stained with peroxidase-conjugated anti-activated caspase-3 antibodies (brown). Original magnification X40. **B,** Representative images of lung tissue from pristane or pristane+GSK treated B6 mice stained with alkaline phosphatase-conjugated anti-activated caspase-3 (red) plus peroxidase-conjugated anti-CD31 (brown, left) or F4/80 (brown, middle and right panels). Original magnification X100. **C,** Immunoperoxidase staining for CD31/Pecam-1 (brown) in blood vessel walls from untreated B6 mice (left), pristane-treated B6 mice (middle), and pristane+GSK treated B6 mice (right). Yellow arrows, staining of small blood vessels with anti-CD31 (immunoperoxidase). Original magnification X100. **D-G,** B6 and BALB/c mice were injected with pristane (Pris). *Pecam1* (CD31) expression was assessed at d-14 by qPCR. **D,** Effect of pristane on *Pecam1* expression in B6 vs. BALB/c lung (qPCR). Pris, pristane-treated; (-), untreated. **E,** Effect of pristane on *Pecam1* expression in lung from B6 vs. B6-*Ccr2*-/- mice (qPCR). **F,** *Pecam1* expression (qPCR) in B6 lung from 0-14 days after pristane treatment. **G,** Flow cytometry of peripheral blood cells for CD31 and CD45. Circulating CD31^+^CD45^-^ cells increased in pristane-treated B6 mice (representative of 3 experiments). * P < 0.05, ** P < 0.01, *** P < 0.001, t-test. ns, not significant. (-), untreated mice.

Consistent with immunohistochemical staining suggesting that pristane may injure lung endothelial cells **(Fig. 2C)**, pristane decreased mRNA expression of *Pecam1* (an endothelial cell-specific marker) in B6, but not BALB/c, lung **(Fig. 2D)**. The effect on *Pecam1* did not rely on monocyte infiltration, since it also was seen in B6-*Ccr2*-/- mice **(Fig. 2E)**. The level of *Pecam1* mRNA decreased with time after pristane treatment **(Fig. 2F)**, consistent with a progressive impairment of endothelial integrity. Coincident with decreased pulmonary CD31 expression, circulating CD31^+^ cells (negative for hematopoietic cell marker CD45) were seen in pristane-treated B6 mice **(Fig. 2G)**, further suggesting that pristane may cause lung endothelial cell injury and shedding of CD31^+^ endothelial cells into the circulation. Alternatively, these cells might represent endothelial cell progenitors released from the bone marrow.

Pristane-treated B6, but not BALB/c, mice develop an ER stress response in the lung (3). Consistent with our prior observations, pristane treatment significantly decreased the expression of *Copa* (consistent with ER/Golgi stress) and increased expression of the ER stress markers *Eif2a* and *Hspa5* (encoding BiP) **(Fig. 3A-C)**. Expression of all three markers returned to normal levels in mice treated with pristane + GSK, suggesting that MEK1/2 inhibition prevents lung injury upstream from the induction of ER stress. Flow cytometry of dissociated lung cells showed increased BiP staining in a subset of CD45^-^ cells from pristane-treated B6 mice **(Fig. 3D)**. We previously reported that these cells express endothelial cell markers (3). Treatment with GSK along with pristane restored BiP staining in the CD45^-^ subset to baseline levels, suggesting that MEK1/2 inhibition blocks the induction of ER stress in endothelial cells. BAL was performed to assess lung inflammation in untreated mice, mice treated with pristane alone, and mice treated with pristane + GSK. Levels of the monocyte-attractive chemokine Ccl2 in the BAL washings (by ELISA) were higher in pristane-treated mice vs. untreated controls and returned to normal in mice treated with pristane + GSK **(Fig. 3E)**, suggesting that MEK1/2 inhibition blocks lung inflammation in pristane-treated mice. GSK treatment significantly reduced annexin V staining of CD31^+^ endothelial cells from pristane-treated B6 mice, indicating that MEK1/2 inhibition decreased the concentration of phosphatidylserine in the outer leaflet of the plasma membrane **(Fig. 3F)**. In view of the reduced numbers of activated caspase-3^+^ apoptotic cells in mice treated with pristane + GSK **(Fig. 2A)**, the reduced annexin V staining of CD31^+^ cells suggests that MEK1/2 inhibition protected endothelial cells from apoptotic cell death.

**Figure 3.**
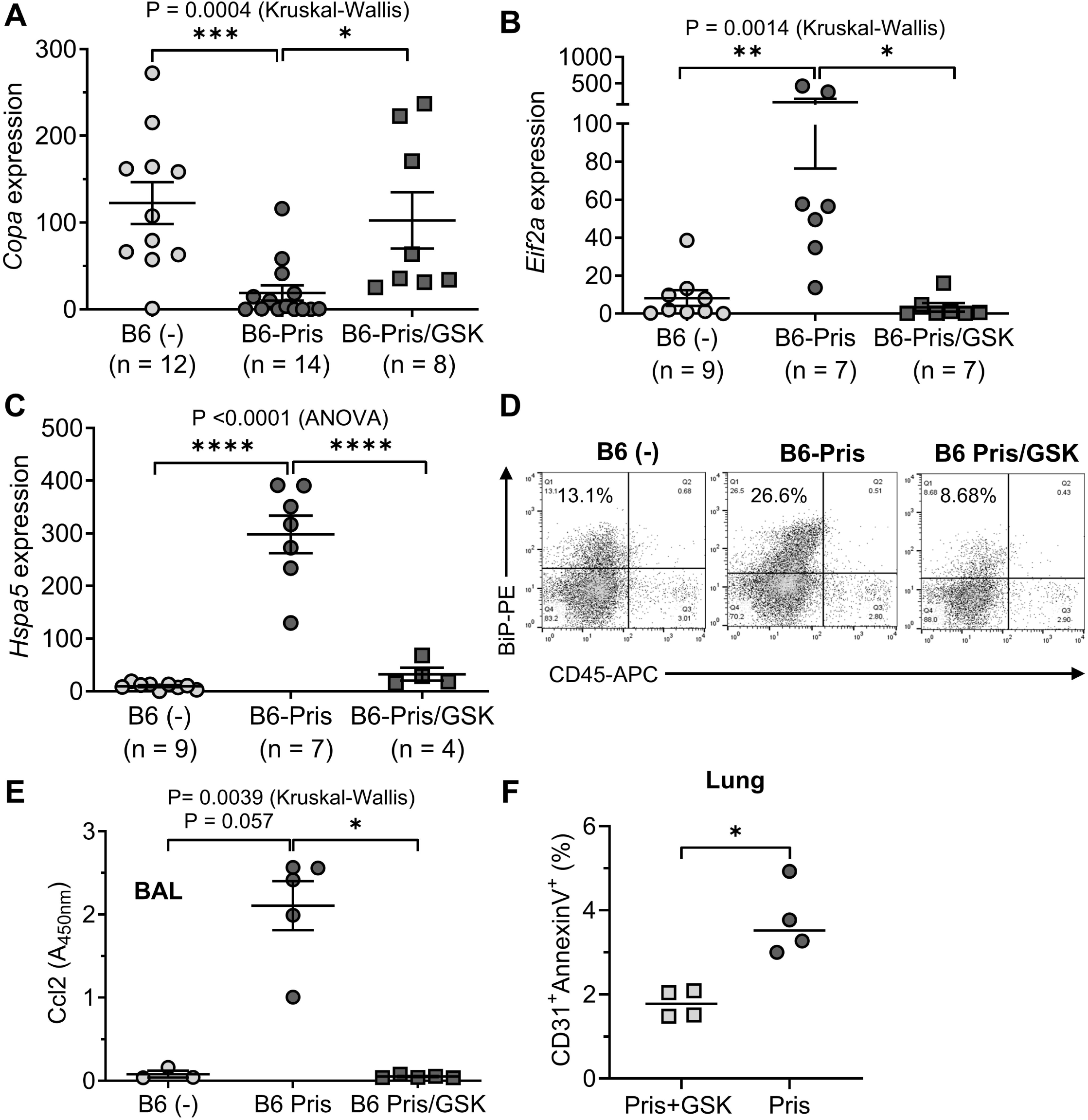
MEK1/2 inhibition alleviates ER stress and lung inflammation. B6 mice were injected i.p. with pristane (Pris) ± daily GSK treatment. 14-d later bronchoalveolar lavage (BAL) was performed and RNA was extracted from lung tissue. Gene expression was determined by qPCR: **A,** *Copa*; **B,** *Eif2a*; **C,** *Hspa5* (BiP). **D,** representative flow cytometry of dissociated lung cells from B6 mice treated with Pris or Pris+GSK vs. untreated (-) mice. **E,** Ccl2 protein in BAL from B6 mice treated with Pris, Pris+GSK, or left untreated (-) was measured by ELISA. * P < 0.05, ** P < 0.01, *** P < 0.001, **** P < 0.0001 (Kruskal Wallis or ANOVA). **F,** B6 mice were treated with pristane ± GSK and at d-6, lung was dissociated and a single cell suspension was analyzed by flow cytometry with anti-CD31 antibodies plus annexin-V staining. * P < 0.05, Mann-Whitney.

The MEK1/2-ERK1/2 pathway controls Egr1, a transcription factor that regulates coagulation, inflammation, vascular permeability, apoptosis, and other pathways (25, 26). Pristane treatment dramatically increased expression of *Egr1* in B6, but not BALB/c, mice, mirroring susceptibility to DAH induction **(Fig. 4A)**. Egr1 also is induced in lung ischemia-perfusion injury (30), and its critical role is supported by the enhanced expression in the vasculature of Egr1-regulated genes mediating ischemic tissue damage, including inflammatory response genes (*Icam1*, *Il1b*, and *Ccl2*), genes involved coagulation (TF/*F3*; PAI-1/*Serpine1*), and *Vegfa*, which is angiogenic but also affects vascular permeability (30). Pristane upregulates expression of the Egr1-regulated gene *Ccl2* (3, 31). Ccl2 protein also increased in BAL from pristane-treated mice **(Fig. 3E)**, but expression of another Egr1-regulated gene *F3* (encoding tissue factor, TF), *decreased* in B6, but not BALB/c, mice **(Fig. 4B)**. TF initiates the extrinsic coagulation cascade and low levels are associated with bleeding (32). *Egr1* induction in B6 mice was blocked by GSK **(Fig. 4C)** and GSK treatment also restored the expression level of F3 back to normal in B6 mice **(Fig. 4D)**.

**Figure 4.**
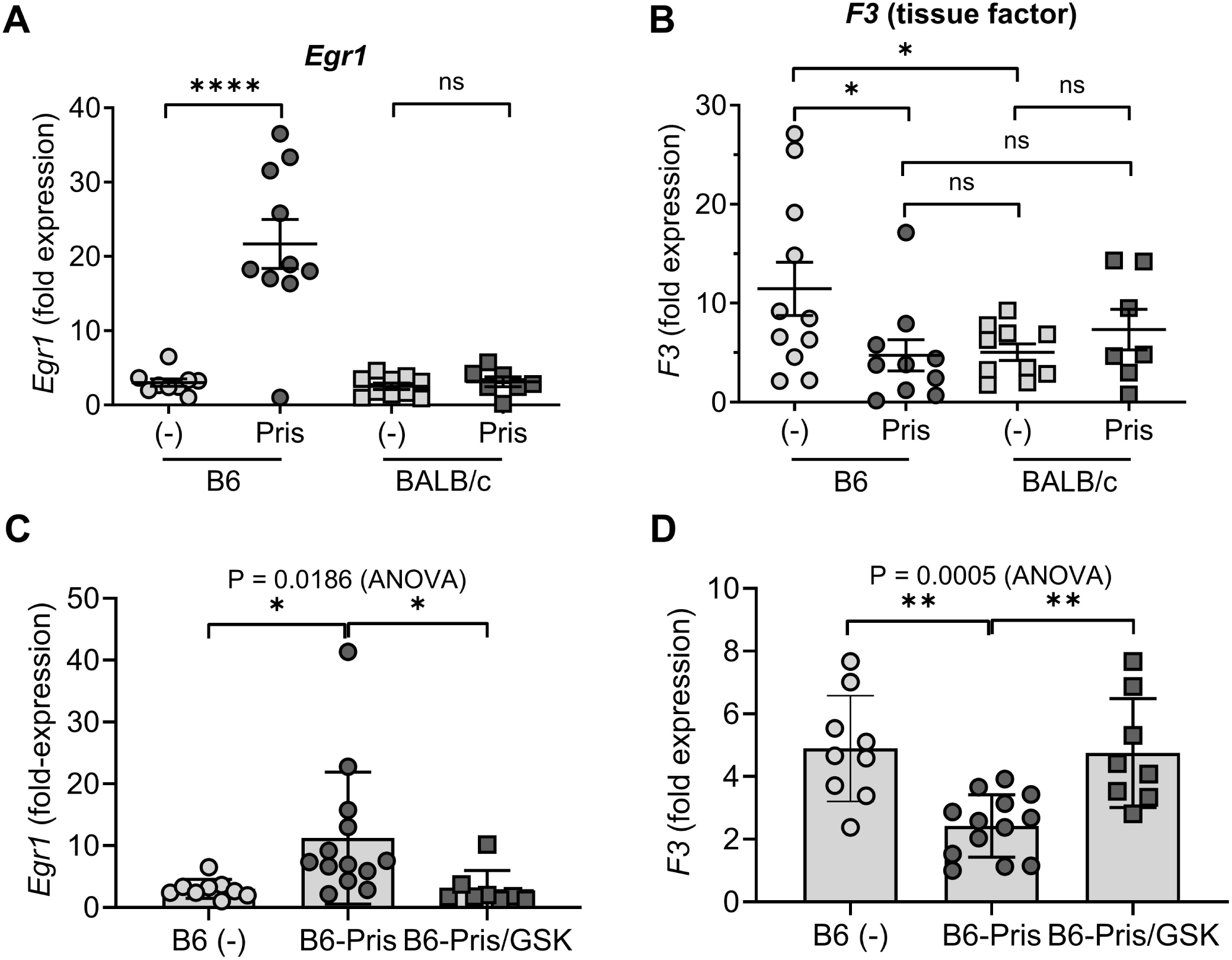
Pristane increases *Egr1* and decreases tissue factor (*F3*) in B6 lung and it is reversed by GSK. **A-B,** B6 and BALB/c mice were treated with pristane ± GSK1120212 (GSK) and expression of *Egr1* **(A)** and *F3* **(B)** was determined in total lung mRNA by qPCR at d-14. *, P < 0.05, **** P < 0.0001 (t-test). ns, not significant. (-) untreated mice; Pris, pristane-treated mice. **C-D,** effect of the MEK1/2 inhibitor GSK on *Egr1* **(C)** and *F3* **(D)** expression in B6 mice. *, P < 0.05, **, P < 0.01 (ANOVA).

The injured endothelium reduces expression of anticoagulant molecules and TF initiates the extrinsic coagulation cascade, generating thrombin **(Fig. 5A)** (33). Conversely, to prevent unnecessary clotting, the uninjured endothelium produces anticoagulants such as tissue factor pathway inhibitor-1 (Tfpi1, henceforth referred to as “Tfpi”) and thrombomodulin (Thbd). Tfpi is the main regulator of TF activity (34) and Thbd is an endothelial cell membrane protein that reduces coagulation by converting thrombin from a procoagulant to an anticoagulant enzyme (33). The balance between pro- and anti-coagulant factors influences the outcome of endothelial cell injury (clotting vs. bleeding) **(Fig. 5A)**.

**Figure 5.**
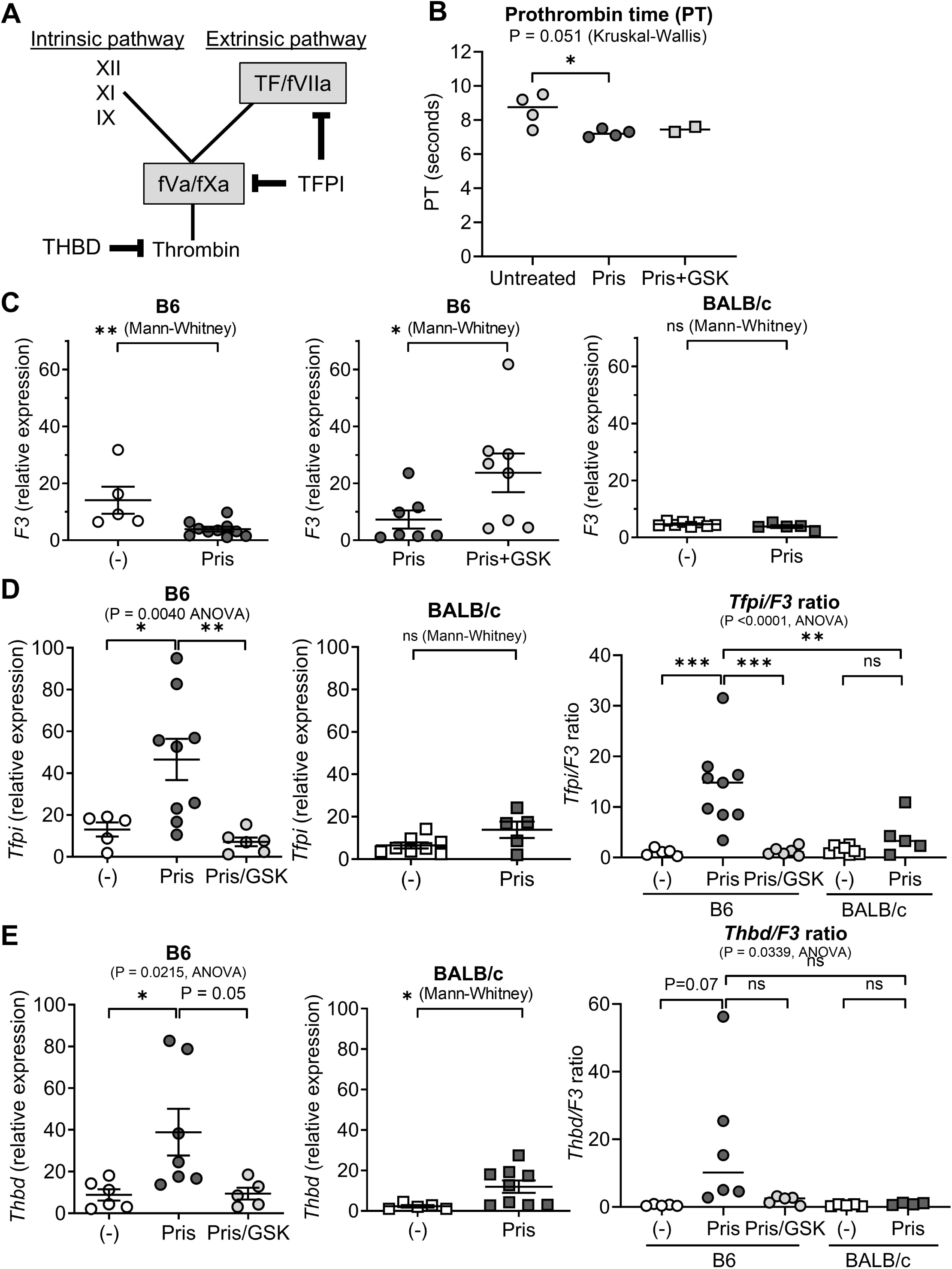
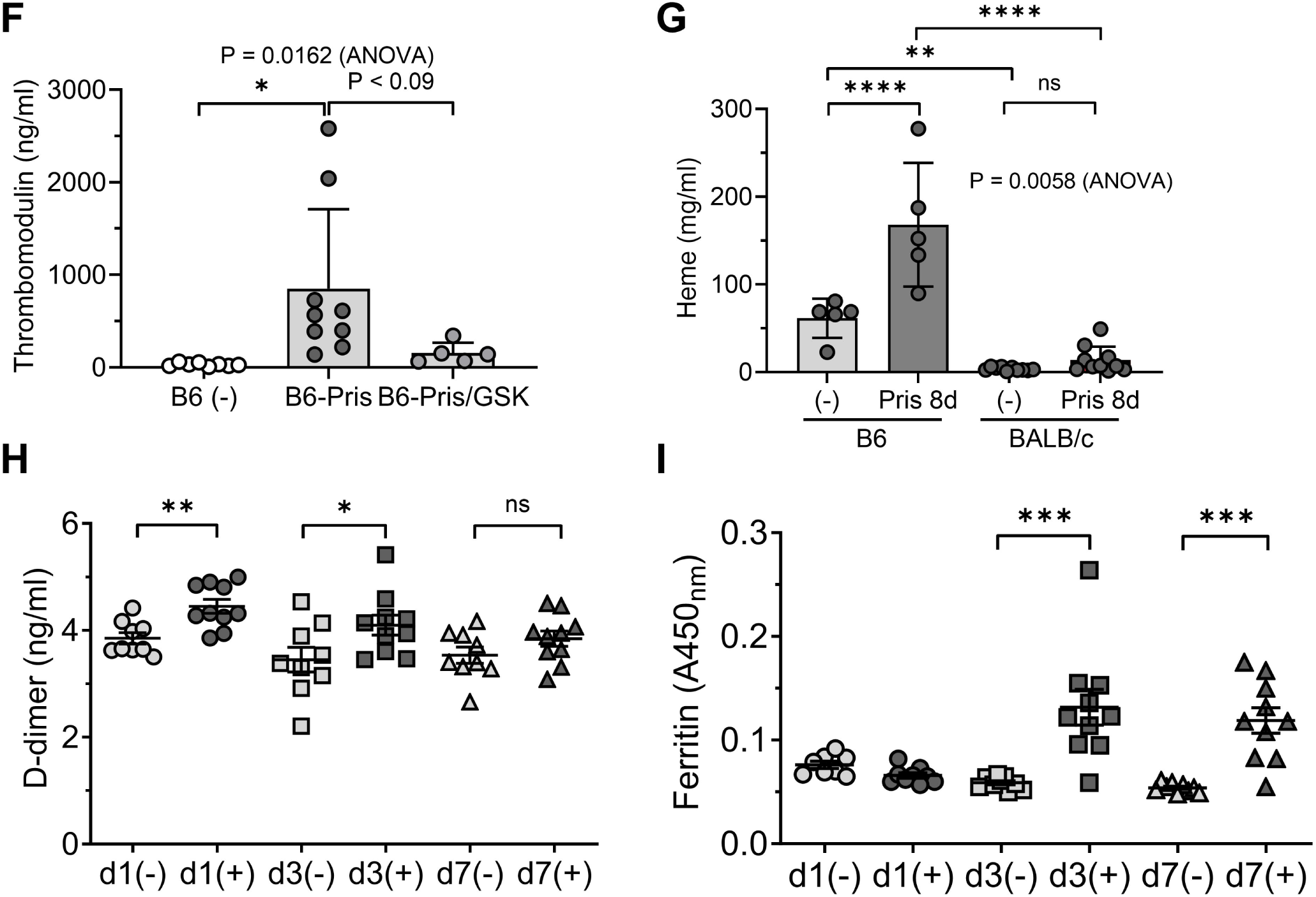
Strain-specific effect of pristane treatment on hemostasis. **A,** Tissue factor pathway inhibitor (TFPI) inhibits clotting via the extrinsic pathway, but clotting can occur via the intrinsic pathway. When TFPI activity is low, tissue factor (TF) generates sufficient thrombin for hemostasis without requiring amplification from the intrinsic pathway. Thrombomodulin (THBD) reduces coagulation by converting thrombin from a procoagulant to an anticoagulant enzyme. **B,** B6 mice were treated with pristane (Pris) or Pris + GSK, or were left untreated. Blood was collected intro citrate tubes by cardiac puncture. PT, INR, and aPTT were determined on an ACL TOP 750 coagulation analyzer. *, P < 0.05 vs. untreated (Kruskal-Wallis with multiple comparisons). **C,** B6 and BALB/c mice were treated with Pris or Pris + GSK120212 (GSK) or left untreated (-). Expression of *F3* (TF) was measured by qPCR at d-10. *, P < 0.05, **, P < 0.01, (Mann-Whitney). ns, not significant. **D-E,** B6 and BALB/c mice were treated with Pris or Pris + GSK or left untreated (-). Expression of *Tfpi* **(D)** and *Thbd* **(E)** was measured at d-10. *Right panels*, ratio of anticoagulant to procoagulant gene expression: *Tfpi/F3* ratio **(C)** *Thbd/F3* ratio **(D)**. * P < 0.05; ** P < 0.01; *** P < 0.001 (ANOVA or Mann-Whitney). Ns, not significant. **F,** plasma thrombomodulin levels (ELISA) in B6 mice treated with Pris, Pris+GSK, or untreated (-). * P < 0.1, ANOVA. **G,** B6 and BALB/c mice were treated with Pris for 8-days or left untreated (-). Tail bleeding was measured as described in the Methods and heme content was quantified by spectrophotometry (A_400 nm_) using a Heme Assay Kit. ** P < 0.01; **** P < 0.0001, ANOVA. **H-I,** B6 mice were injected i.p. with pristane (+) or were untreated (-). Serum D-dimer **(H)** and ferritin **(I)** levels were measured by ELISA at days 1, 3, and 7 after pristane treatment. * P < 0.05; **, P < 0.01; *** P < 0.001; (t-test). ns, not significant

Although the PT is a measure of overall extrinsic pathway activity, it does not adequately reflect the balance of TF and its inhibitors on the endothelium. For example, a monoclonal antibody that blocks the interaction of TFPI with TF promotes coagulation and hemostasis following tail injury in hemophilia mice by increasing extrinsic pathway activity, but does not alter the PT (or aPTT) (35). Consistent with those observations, the PT **(Fig. 5B)** and INR (not shown) was not prolonged in pristane-treated mice, and in fact was shortened modestly. GSK therapy had little effect on the PT. Similarly, the aPTT (a measure of intrinsic pathway activity) was shortened slightly by pristane and unaffected by GSK (not shown).

In untreated mice, *F3* (TF) expression was higher in B6 vs. BALB/c mice (P < 0.002, Mann Whitney) (not shown, compare **Fig. 5C** left vs. far right). Pristane injection decreased *F3* expression in B6 mice, but F3 was expressed at similar levels in pristane-treated vs. untreated BALB/c mice. GSK treatment restored *F3* expression to the baseline (untreated) level in B6 mice. *Tfpi* expression increased in pristane-treated mice and concomitant administration of GSK restored it to baseline levels **(Fig. 5D)**. In contrast, pristane treatment had little effect on *Tfpi* expression in BALB/c mice. The expression of *Tfpi* was higher in pristane-treated B6 vs.

BALB/c mice (P = 0.04, Mann-Whitney, not shown). The ratio of *Tfpi*/*F3*, which may provide a measure of the relative predisposition of the endothelium to bleeding vs. hemostasis, was similar in untreated B6 vs. BALB/c mice **(Fig. 5D)**. However, after pristane treatment, the *Tfpi*/*F3* ratio increased markedly in B6 mice with a less pronounced (and not statistically significant) trend in BALB/c **(Fig. 5D)**. The *Tfpi*/*F3* ratio was significantly higher in pristane-treated B6 vs. BALB/c mice. Treating B6 mice with GSK along with pristane restored the *Tfpi*/*F3* ratio to its baseline level.

*Thbd* expression exhibited a similar pattern **(Fig. 5E)**. Pristane treatment increased *Thbd* expression in both B6 and BALB/c mice, and the expression was normalized in B6 mice by concomitant GSK treatment. *Thbd* expression was higher in pristane-treated B6 vs. BALB/c mice (P = 0.04, Mann-Whitney, not shown). As in the case of the *Tfpi/F3* ratio, there was a trend toward an increased *Thbd/F3* ratio in pristane-treated B6 mice vs. untreated controls that was not apparent in BALB/c mice **(Fig. 5E)**.

Increased *Thbd* expression was associated with an increased level of soluble thrombomodulin (sThbd) protein detected by ELISA in the plasma of pristane-treated mice **(Fig. 5F)**. GSK therapy reduced the level of plasma sThbd in pristane-treated mice, though it did not reach statistical significance. Together, these studies suggest that due to shifts in the relative levels of TF (F3) and its inhibitors (Tfpi and Thbd), pristane treatment may promote an anticoagulant state in the B6 lung endothelium that is absent in the endothelium of pristane-treated BALB/c mice. Additionally, the changes induced by pristane were reversed by GSK treatment, suggesting that the relative anticoagulant state induced by pristane was dependent on the MEK1/2 pathway.

Hemostasis in pristane-treated mice was assessed at a functional level using a tail bleeding assay that is dependent on the extrinsic coagulation pathway (35). This assay was used previously to show genetic differences in hemostasis between B6 and BALB/c mice that correlate with circulating Tfpi activity (21). We measured tail bleeding in untreated B6 and BALB/c mice vs. pristane-treated mice 8-days after pristane treatment (prior to the onset of alveolar hemorrhage in pristane-treated B6 mice). This timepoint was selected because DAH causes substantial blood loss/anemia at 14-d in B6 mice (H Zhuang, unpublished data). Tail bleeding was higher in untreated B6 vs. BALB/c mice, confirming previous results (21) **(Fig. 5G)**. In addition, tail bleeding increased markedly in pristane-treated B6 mice vs. untreated controls **(Fig. 5G)**. There was a small, but not statistically significant, trend in BALB/c mice. Tail bleeding was substantially higher in pristane-treated B6 vs. BALB/c mice.

Pristane treatment induces pulmonary microvascular injury (3). Since sThbd released from injured endothelial cells is a marker for endothelial cell injury in patients with COVID infection (36), the increased circulating Thbd in pristane-treated mice **(Fig. 5F)** provides further evidence of endothelial injury. Endothelial injury in COVID infection causes microthrombi and increased D-dimer levels (37). Similarly, serum D-dimer increased at days 1 and 3 after pristane treatment, returning to baseline at day-7 **(Fig. 5H)**. COVID infection also causes high serum ferritin levels (38), consistent with its association with ML activation (39). As pristane-induced DAH is monocyte/ML dependent, we measured serum ferritin in pristane-treated mice **(Fig. 5I)**. Despite the increased D-dimer, ferritin was normal at day-1 after pristane treatment. However, the ferritin was significantly higher at days 3 and 7. Thus, there are some similarities between human COVID infection, which causes lung microvascular injury, and pristane-induced DAH in mice. However, unlike the situation in COVID infection, pristane may cause transient thrombosis (days 1-3), which is limited by vigorous Tfpi/Thbd production.

## Discussion

Pristane-induced lupus is associated with DAH in B6 and B10 mice, but not other strains, such as BALB/c (1). Damage to the lung microvascular endothelium resulting in ER stress and cell death plays a central role in the pathogenesis of DAH (3). The resistance of BALB/c mice to DAH can be overcome by treating with a low dose of the ER stress inducer thapsigargin along with pristane, suggesting that the severity of endothelial damage is one determinant of whether or not DAH develops. However, the inflammatory pathway(s) mediating DAH and the existence of other factors governing susceptibility to DAH remain poorly understood. The present study suggests that activation of the MEK1/2-ERK1/2 pathway by pristane is involved in microvascular injury, causing an anti-thrombotic state by altering the balance of TF (*F3*) and its main regulator, Tfpi. Inhibition of either MEK1/2 (GSK) or ERK1/2 (SCH) abolished DAH, prevented endothelial injury as indicated by prevention of ER stress, decreased CD31 (PECAM) expression and staining, and increased activated caspase (apoptosis) staining. GSK therapy also normalized the hemostasis defect unique to pristane-treated B6 mice.

### Impaired hemostasis is a risk factor for DAH

TF is a transmembrane protein expressed by perivascular cells that binds factor VIIa (40). Following endothelial injury, the TF-factor VIIa complex proteolytically activates factors IX and X, ultimately leading to the formation of a fibrin clot **(Fig. 5A)**. To prevent unnecessary clotting, the uninjured endothelium produces anticoagulants such as Tfpi and Thbd. Thbd is an endothelial cell membrane protein that reduces coagulation by converting thrombin from a procoagulant to an anticoagulant enzyme (33).

Deficiency causes a bleeding disorder (41). Tfpi is a serine protease inhibitor produced by endothelial cells and megakaryocytes that is the main regulator of TF activity (34). The balance between pro- and anti-coagulant factors influences the outcome of endothelial cell injury (clotting vs. bleeding). Inhibition of Tfpi corrects the hemostasis defect in animal models of hemophilia and a Tfpi neutralizing antibody (concizumab) is undergoing testing in human hemophilia (42, 43).

The risk of DAH increases in hemophilia patients (odds ratio 7.47) (6, 44, 45). Low levels of TF also increase the risk. In mice with low TF, intratracheal LPS or influenza A virus infection causes DAH (46, 47). In humans, impaired TF expression promotes DAH in influenza pneumonia (48) and heterozygous TF deficiency causes a bleeding disorder (32). Human DAH has been treated successfully with recombinant factor VII (rFVII), which binds TF and is activated to FVIIa, or with rFVIIa (49, 50). DAH also can be induced by influenza infection in mice treated with warfarin or the thrombin inhibitor dabigatran etexilate (51) and is more prevalent in patients receiving warfarin or aspirin (52).

Our studies suggest that the balance between TF and its inhibitors was disrupted by pristane treatment, as indicated by an increased *Tfpi/F3* ratio and increased sThbd in the plasma **(Fig. 5D, F)**. These abnormalities were corrected by GSK treatment. The increased *Tfpi*/*F3* ratio was mainly due to increased *Tfpi* expression. A prior study shows that at baseline, B6 mice are more prone to bleed than BALB/c and that B6 mice have high levels of circulating Tfpi (21). However, increased tail bleeding in untreated B6 vs. BALB/c mice is not reflected by *ex vivo* coagulation assays such as the PT or aPTT (53). We confirmed the mild bleeding tendency in B6 mice and uncovered a more pronounced interstrain difference after pristane treatment **(Fig. 5)**.

Tail bleeding more than doubled 8-d after treating B6 mice with pristane, whereas it did not increase significantly in BALB/c mice **(Fig. 5G)**. Our data suggest that the increased levels of Tfpi and/or circulating Thbd contribute to the high susceptibility of B6 vs. BALB/c mice to pristane-induced DAH, implicating dysregulation of TF in the pathogenesis of DAH.

Pristane treatment injures lung microvascular endothelial cells (3) and the present studies indicate that it impairs the expression of Pecam-1 (CD31) **(Fig. 2)**, a protein that maintains endothelial integrity (28, 29). We previously reported that pristane treatment increases annexin-V staining of lung endothelial cells, consistent with higher levels of phosphatidylserine in the outer leaflet of the plasma membrane (3). MEK1/2 inhibition reduced annexin V staining of CD31^+^ lung endothelial cells **(Fig. 3F)** and inhibited lung inflammation as indicated by the normalization of Ccl2 in BAL **(Fig. 3E)**. This is significant because increased exposure of phosphatidylserine on the endothelial cell surface promotes both thrombosis and inflammation in COVID infection (54). In addition, phosphatidylserine exposure on apoptotic cells results in complement activation, coating of the cell surface with C3b, and recognition by phagocytes bearing CR3 (CD11b/CD18) and CR4 (CD11c/CD18) (55). B6 mice deficient in either C3 or CD18 are not susceptible to pristane-induced DAH (2).

In COVID infection, exposure of phosphatidylserine promotes the formation of TF-FVIIa-FX complexes, leading to activation of the extrinsic pathway and a prothrombotic state (56). We found evidence of thrombosis (elevated D-dimer) as early as 1-d after pristane treatment **(Fig. 5H)**, consistent with the idea that endothelial injury and transient thrombosis precedes DAH. TF is exposed to the plasma only after endothelial injury, initiating coagulation, whereas excessive clot formation is prevented by inhibition of TF (Tfpi) and thrombin (Thbd) (57). Pristane treatment considerably increased both inhibitors, suggesting that the downregulation of this early prothrombotic state may be inappropriately vigorous.

TF activates factor VII (the extrinsic pathway), and TF-FVIIa converts FX to FXa, activating the common pathway (factors X, V, II/prothrombin, and fibrinogen). The prothrombin time (PT) assesses activation of the extrinsic and common pathways (57). However, neither the PT nor the activated partial thromboplastin time (aPTT, a measure of activation of the intrinsic and common pathways) is affected by TF activity, as illustrated by the observation that a *TFPI* variant (T-287C) associated with increased risk of thrombosis, coronary artery disease, and high plasma fibrinogen levels does not alter the PT, aPTT, or thrombin time (58). Thus, it is not surprising that the PT and aPTT were not prolonged in pristane treated vs. untreated B6 mice (**Fig. 5B** and data not shown) since the PT and aPTT are of limited value for evaluating TF activity. However, we cannot exclude the possibility that pristane might cause subtle abnormalities of the intrinsic/extrinsic coagulation cascade or that it impairs platelet function in B6 mice. Further studies are needed to examine this possibility.

Interestingly, although increased tail bleeding in B6 mice might suggest a systemic bleeding tendency **(Fig. 5)**, pristane-treated B6 mice only have lung hemorrhage, and do not bleed into other tissues such as the kidney or brain (H. Zhuang, unpublished observation). Bleeding and thrombotic disorders often show a predilection for specific sites, suggesting an important role for local factors (59). As in pristane-induced lupus-DHA, SLE patients with DAH do not develop skin, mesenteric, renal, or nervous system vasculitis or hemorrhage. This selectivity most likely reflects the marked heterogeneity of microvascular endothelial cells from different tissues (60). The substantial differences in gene expression between microvascular endothelial cells from lung vs. other tissues (61) could influence susceptibility to endothelial injury, which may then initiate activation of the coagulation cascade. Also, regulators of hemostasis, including Thbd, are not uniformly distributed in the vasculature (59). Endothelial gene expression disparities between tissues may explain why hemorrhage in pristane-treated mice and SLE patients is restricted to the lung. Polymorphisms of the differentially expressed genes may partly underlie the susceptibility of B6 vs. BALB/c lung to pristane-induced lung injury.

### DAH susceptibility is regulated by the MEK1/2-ERK1/2 pathway

Endothelial cell injury, ER stress, cell death, and inflammation are seen consistently in lung from B6 mice with DAH, whereas pristane did not induce any of these abnormalities in BALB/c mice and tail bleeding was not augmented **[Figs. 1, 5** and ref. (3)]. In B6 mice, endothelial injury and cell death **(Figs. 2-3)**, ER stress **(Fig. 3)**, and inflammation **(Figs. 3E and 4)** all were blocked by GSK therapy. A second MEK1/2 inhibitor (AZD8330) also was protective and two different ERK1/2 inhibitors (SCH and ulixertinib) prevented DAH **(Fig. 1)**. *F3*, *Tfpi*, and *Thbd* expression and plasma sThbd returned to normal in B6 mice receiving GSK and the balance between F3 and its inhibitors was normalized **(Fig. 5)**. In contrast, inhibition of other MAP kinases (JNK, p38) and activation of cAMP signaling had no effect on DAH, suggesting that the MEK1/2-ERK1/2 pathway is uniquely involved in the pathogenesis of DAH **(Fig. 1H)**. [flow cytometry-phosphoERK?]

Consistent with its regulation by MEK1/2-ERK1/2, pristane dramatically increased *Egr1* expression in B6, but not BALB/c, mice **(Fig. 4)**. Egr1 is a transcription factor downstream of MEK1/2 that regulates coagulation, inflammation, vascular permeability, and apoptosis (25, 26) and plays a central role in lung ischemia-perfusion injury (30). Pristane upregulates the Egr1-regulated gene *Ccl2* (3, 31) and GSK therapy prevented pristane-induced *Egr1* induction and normalized production of Ccl2 in the lung **(Fig. 3E)**. Egr1 also regulates *Serpine1* (endothelial plasminogen activator inhibitor-1, the main inhibitor of tissue plasminogen activator), but the frequency of DAH is comparable to controls in *Serpine1*-deficient mice (62). Paradoxically, although Egr1 upregulates *F3*, pristane treatment decreased *F3* expression and GSK therapy normalized it in B6 mice **(Fig. 5)**. In contrast, upregulation of *Tfpi* and *Thbd* by pristane was prevented by GSK, although there is little evidence that these genes are MEK1/2-ERK1/2-regulated (63). An alternative explanation is that inhibition of the MEK1/2-ERK1/2 pathway might prevent DAH by normalizing expression of the liver X receptor (LXR)-regulated reverse cholesterol transporter ATP-binding cassette subfamily A member 1 (Abca1). Abca1 expression decreases in pristane-treated B6 mice and is normalized by LXR agonist therapy, preventing DAH (24). ERK1/2 activation decreases Abca1 expression (64), so GSK and SCH might act in a manner similar to that of the LXR agonist T0901317 (24).

Regardless of the mechanism(s) involved, the prevention of endothelial injury, ER stress, and apoptosis by GSK therapy **(Figs. 2-3)** suggests that activation of the MEK1/2-ERK1/2 pathway is an early event in the pathogenesis of DAH. Although ERK1/2 activation typically promotes cell proliferation and survival, it also can be pro-apoptotic (65). Our working model is that pristane-induced damage to the lung microvascular endothelium is MEK1/2-ERK1/2-dependent, resulting in exposure of phosphatidylserine on the endothelial cell membrane, which promotes an initial TF-mediated procoagulant state followed by the inappropriately high production of anticoagulant proteins (Tfpi and Thdb), causing alveolar hemorrhage. Further studies will be needed to validate this model.

### Clinical implications

DAH is associated with lupus and other rheumatological autoimmune disorders (6). Although severe alveolar hemorrhage is seen in only 3-4% of SLE patients, subclinical focal hemorrhage is seen at autopsy in the lungs of 30-66% of patients (*5*). DAH with pulmonary capillaritis also may be seen in rheumatoid arthritis (66), primary and secondary antiphospholipid syndrome in the absence of thrombosis (67, 68), and patients treated with certain medications including anticoagulants and mechanistic target of rapamycin (mTOR) inhibitors such as sirolimus (rapamycin) and everolimus (69, 70). As in SLE, DAH in these conditions is more prevalent in women than men (6). However, it is not known why severe DAH affects only 3-4% of lupus patients.

Our data suggest that, as in B6 vs. BALB/c mice, subtle hereditary differences in regulating the extrinsic coagulation pathway may influence susceptibility to DAH. There is a group of otherwise healthy individuals with “bleeding of unknown cause” (BUC) and normal hemostasis tests (71). Most lack a diagnosis. Next generation and whole genome sequencing strategies help, but only 3.2% of patients with unexplained bleeding and normal hemostasis tests can be assigned a molecular diagnosis (72). Although BUC is not rare, evaluation is challenging (71). Even in von Willebrand disease, plasma vWF levels correlate only weakly with bleeding. BUC is diagnosed by determining a “bleeding score” (71). The PT and aPTT are slightly prolonged, but remain within the normal ranges and are not helpful diagnostically. The most reliable lab abnormality in patients with recurrent epistaxis, bruising, or menorrhagia typical of BUC may be an increased soluble TFPI (sTFPI) protein level (73). Like patients with BUC, B6 mice have a subtle bleeding defect that is evaluated clinically by measuring the tail bleeding and is associated with high sTFPI (21). The PT and aPTT are similar in B6 vs. BALB/c mice (53). We confirmed the mild bleeding disorder in untreated B6 mice and found that it was aggravated by pristane treatment **(Fig. 5G)**, in association with increased sThbd in the blood **(Fig. 5F)** and increased *Thbd* mRNA in lung **(Fig. 5E)**. Besides inhibiting the extrinsic coagulation pathway, sThbd is a marker of EC injury and may be a biomarker for cardiovascular disease in SLE (74, 75).

Our studies raise the possibility that inflammation/endothelial injury could promote bleeding by increasing TFPI and/or THBD. However, it remains to be determined whether dysregulation of TFPI/THBD plays a causal role in DAH and the relative importance of each factor. Tools are available to identify individuals with bleeding tendencies (76, 77). It may be of interest to investigate whether screening would distinguish the 3-4% of SLE patients who are susceptible to life-threating DAH from those with subclinical alveolar hemorrhage (30-66%) or no hemorrhage at all. If MEK1/2-regulated TFPI (or THBD) expression does play a causal role in DAH, there may be a potential to treat alveolar hemorrhage with neutralizing antibodies against TFPI (concizumab) (42, 43) or with FDA-approved MEK1/2 inhibitors, such as trametinib (GSK1120212) (27).

## Acknowledgments

We thank the University of Florida Molecular Pathology Core for performing histology and immunohistochemistry of lung tissues. We thank Dr. William L. Clapp (Director of Renal Pathology, Department of Pathology, Immunology and Laboratory Medicine, University of Florida) for reviewing slides of renal tissue from B6 mice and Ms. Heather D. Pruitt for carrying out the coagulation (PT and aPTT) assays.

## References

1. Barker TT, Lee PY, Kelly-Scumpia KM, Weinstein JS, Nacionales DC, Kumagai Y, et al. Pathogenic role of B cells in the development of diffuse alveolar hemorrhage induced by pristane. LabInvest. 2011;91:1540–50.

2. Zhuang H, Han S, Lee PY, Khaybullin R, Shumyak S, Lu L, et al. Pathogenesis of diffuse alveolar hemorrhage in murine lupus. Arthritis Rheumatol. 2017;69:1280–93 (see commentary).

3. Zhuang H, Hudson E, Han S, Arja RD, Hui W, Lu L, et al. Microvascular lung injury and endoplasmic reticulum stress in SLE-associated alveolar hemorrhage and pulmonary vasculitis. AmJPhysiolLung CellMolPhysiol. 2022;323(6):L715–L29.

4. Tumurkhuu G, Laguna DE, Moore RE, Contreras J, Santos GL, Akaveka L, et al. Neutrophils Contribute to ER Stress in Lung Epithelial Cells in the Pristane-Induced Diffuse Alveolar Hemorrhage Mouse Model. FrontImmunol. 2022;13:790043.

5. Al-Adhoubi NK, Bystrom J. Systemic lupus erythematosus and diffuse alveolar hemorrhage, etiology and novel treatment strategies. Lupus. 2020;29(4):355–63.

6. Kambhatla S, Vipparthy S, Manadan AM. Rheumatic diseases associated with alveolar hemorrhage: analysis of the national inpatient sample. ClinRheumatol. 2023;42(4):1177–83.

7. Balcan B, Simsek E, Ugurlu AO, Demiralay E, Sahin S. Sirolimus-Induced Diffuse Alveolar Hemorrhage: A Case Report. AmJTher. 2016;23(6):e1938–e41.

8. Lee PY, Kumagai Y, Li Y, Takeuchi O, Yoshida H, Weinstein J, et al. TLR7-dependent and FcgammaR-independent production of type I interferon in experimental mouse lupus. JExpMed. 2008;205(13):2995–3006.

9. Nacionales DC, Kelly-Scumpia KM, Lee PY, Weinstein JS, Sobel E, Satoh M, et al. Deficiency of the Type I interferon receptor protects mice from experimental lupus. Arthritis Rheum. 2007;56:3770–83.

10. Arthur JS, Ley SC. Mitogen-activated protein kinases in innate immunity. NatRevImmunol. 2013;13(9):679–92.

11. Morris EJ, Jha S, Restaino CR, Dayananth P, Zhu H, Cooper A, et al. Discovery of a novel ERK inhibitor with activity in models of acquired resistance to BRAF and MEK inhibitors. Cancer Discov. 2013;3(7):742–50.

12. Chaikuad A, Tacconi EM, Zimmer J, Liang Y, Gray NS, Tarsounas M, et al. A unique inhibitor binding site in ERK1/2 is associated with slow binding kinetics. NatChemBiol. 2014;10(10):853–60.

13. Gilmartin AG, Bleam MR, Groy A, Moss KG, Minthorn EA, Kulkarni SG, et al. GSK1120212 (JTP-74057) is an inhibitor of MEK activity and activation with favorable pharmacokinetic properties for sustained in vivo pathway inhibition. ClinCancer Res. 2011;17(5):989–1000.

14. Yamaguchi T, Kakefuda R, Tajima N, Sowa Y, Sakai T. Antitumor activities of JTP-74057 (GSK1120212), a novel MEK1/2 inhibitor, on colorectal cancer cell lines in vitro and in vivo. IntJOncol. 2011;39(1):23–31.

15. Akinleye A, Furqan M, Mukhi N, Ravella P, Liu D. MEK and the inhibitors: from bench to bedside. JHematolOncol. 2013;6:27.

16. Roskoski R, Jr. Targeting ERK1/2 protein-serine/threonine kinases in human cancers. PharmacolRes. 2019;142:151–68.

17. Campbell RM, Anderson BD, Brooks NA, Brooks HB, Chan EM, De Dios A, et al. Characterization of LY2228820 dimesylate, a potent and selective inhibitor of p38 MAPK with antitumor activity. MolCancer Ther. 2014;13(2):364–74.

18. Bennett BL, Sasaki DT, Murray BW, O’Leary EC, Sakata ST, Xu W, et al. SP600125, an anthrapyrazolone inhibitor of Jun N-terminal kinase. ProcNatlAcadSciUSA. 2001;98(24):13681–6.

19. Schwede F, Maronde E, Genieser H, Jastorff B. Cyclic nucleotide analogs as biochemical tools and prospective drugs. PharmacolTher. 2000;87(2-3):199–226.

20. Zhuang H, Han S, Xu Y, Li Y, Wang H, Yang LJ, et al. Toll-like receptor 7-stimulated tumor necrosis factor alpha causes bone marrow damage in systemic lupus erythematosus. Arthritis Rheumatol. 2014;66(1):140–51.

21. White TA, Pan S, Witt TA, Simari RD. Murine strain differences in hemostasis and thrombosis and tissue factor pathway inhibitor. ThrombRes. 2010;125(1):84–9.

22. De Pablo-Moreno JA, Liras A, Revuelta L. Standardization of Coagulation Factor V Reference Intervals, Prothrombin Time, and Activated Partial Thromboplastin Time in Mice for Use in Factor V Deficiency Pathological Models. FrontVetSci. 2022;9:846216.

23. Peters LL, Cheever EM, Ellis HR, Magnani PA, Svenson KL, Von Smith R, et al. Large-scale, high-throughput screening for coagulation and hematologic phenotypes in mice. PhysiolGenomics. 2002;11(3):185–93.

24. Han S, Zhuang H, Shumyak S, Wu J, Xie C, Li H, et al. Liver X Receptor Agonist Therapy Prevents Diffuse Alveolar Hemorrhage in Murine Lupus by Repolarizing Macrophages. FrontImmunol. 2018;9:135.

25. Caunt CJ, Sale MJ, Smith PD, Cook SJ. MEK1 and MEK2 inhibitors and cancer therapy: the long and winding road. NatRevCancer. 2015;15(10):577–92.

26. Guha M, O’Connell MA, Pawlinski R, Hollis A, McGovern P, Yan SF, et al. Lipopolysaccharide activation of the MEK-ERK1/2 pathway in human monocytic cells mediates tissue factor and tumor necrosis factor alpha expression by inducing Elk-1 phosphorylation and Egr-1 expression. Blood. 2001;98(5):1429–39.

27. Zeiser R, Andrlova H, Meiss F. Trametinib (GSK1120212). Recent Results Cancer Res. 2018;211:91–100.

28. Cheung KCP, Fanti S, Mauro C, Wang G, Nair AS, Fu H, et al. Preservation of microvascular barrier function requires CD31 receptor-induced metabolic reprogramming. NatCommun. 2020;11(1):3595.

29. Lertkiatmongkol P, Liao D, Mei H, Hu Y, Newman PJ. Endothelial functions of platelet/endothelial cell adhesion molecule-1 (CD31). CurrOpinHematol. 2016;23(3):253–9.

30. Yan SF, Fujita T, Lu J, Okada K, Shan Zou Y, Mackman N, et al. Egr-1, a master switch coordinating upregulation of divergent gene families underlying ischemic stress. NatMed. 2000;6(12):1355–61.

31. Han S, Zhuang H, Arja RD, Reeves WH. A novel monocyte differentiation pattern in pristane-induced lupus with diffuse alveolar hemorrhage. eLife. 2022;11.

32. Schulman S, El-Darzi E, Florido MH, Friesen M, Merrill-Skoloff G, Brake MA, et al. A coagulation defect arising from heterozygous premature termination of tissue factor. JClinInvest. 2020;130(10):5302–12.

33. Watanabe-Kusunoki K, Nakazawa D, Ishizu A, Atsumi T. Thrombomodulin as a Physiological Modulator of Intravascular Injury. FrontImmunol. 2020;11:575890.

34. Wood JP, Ellery PE, Maroney SA, Mast AE. Biology of tissue factor pathway inhibitor. Blood. 2014;123(19):2934–43.

35. Cardinal M, Kantaridis C, Zhu T, Sun P, Pittman DD, Murphy JE, et al. A first-in-human study of the safety, tolerability, pharmacokinetics and pharmacodynamics of PF-06741086, an anti-tissue factor pathway inhibitor mAb, in healthy volunteers. JThrombHaemost. 2018;16(9):1722–31.

36. Boron M, Hauzer-Martin T, Keil J, Sun XL. Circulating Thrombomodulin: Release Mechanisms, Measurements, and Levels in Diseases and Medical Procedures. TH Open. 2022;6(3):e194–e212.

37. Thachil J, Favaloro EJ, Lippi G. D-dimers-“Normal” Levels versus Elevated Levels Due to a Range of Conditions, Including “D-dimeritis,” Inflammation, Thromboembolism, Disseminated Intravascular Coagulation, and COVID-19. SeminThrombHemost. 2022;48(6):672–9.

38. Kurian SJ, Mathews SP, Paul A, Viswam SK, Kaniyoor Nagri S, Miraj SS, et al. Association of serum ferritin with severity and clinical outcome in COVID-19 patients: An observational study in a tertiary healthcare facility. ClinEpidemiolGlobHealth. 2023;21:101295.

39. Kernan KF, Carcillo JA. Hyperferritinemia and inflammation. IntImmunol. 2017;29(9):401–9.

40. Grover SP, Mackman N. Tissue Factor: An Essential Mediator of Hemostasis and Trigger of Thrombosis. ArteriosclerThrombVascBiol. 2018;38(4):709–25.

41. Langdown J, Luddington RJ, Huntington JA, Baglin TP. A hereditary bleeding disorder resulting from a premature stop codon in thrombomodulin (p.Cys537Stop). Blood. 2014;124(12):1951–6.

42. Chowdary P. Inhibition of Tissue Factor Pathway Inhibitor (TFPI) as a Treatment for Haemophilia: Rationale with Focus on Concizumab. Drugs. 2018;78(9):881–90.

43. Matsushita T, Shapiro A, Abraham A, Angchaisuksiri P, Castaman G, Cepo K, et al. Phase 3 Trial of Concizumab in Hemophilia with Inhibitors. NEnglJMed. 2023;389(9):783–94.

44. Chandrakala S, Jijina F, Ghosh K. Diffuse alveolar haemorrhage with severe haemophilia. Haemophilia. 2010;16(6):962–4.

45. Kasai H, Terada J, Hoshi H, Urushibara T, Kato F, Nishimura R, et al. Repeated Diffuse Alveolar Hemorrhage in a Patient with Hemophilia B. InternMed. 2017;56(4):425–8.

46. Bastarache JA, Sebag SC, Clune JK, Grove BS, Lawson WE, Janz DR, et al. Low levels of tissue factor lead to alveolar haemorrhage, potentiating murine acute lung injury and oxidative stress. Thorax. 2012;67(12):1032–9.

47. Antoniak S, Tatsumi K, Hisada Y, Milner JJ, Neidich SD, Shaver CM, et al. Tissue factor deficiency increases alveolar hemorrhage and death in influenza A virus-infected mice. JThrombHaemost. 2016;14(6):1238–48.

48. von Ranke FM, Zanetti G, Hochhegger B, Marchiori E. Infectious diseases causing diffuse alveolar hemorrhage in immunocompetent patients: a state-of-the-art review. Lung. 2013;191(1):9–18.

49. Pathak V, Kuhn J, Gabriel D, Barrow J, Jennette JC, Henke DC. Use of Activated Factor VII in Patients with Diffuse Alveolar Hemorrhage: A 10 Years Institutional Experience. Lung. 2015;193(3):375–9.

50. Alabed IB. Treatment of diffuse alveolar hemorrhage in systemic lupus erythematosus patient with local pulmonary administration of factor VIIa (rFVIIa): a case report. Medicine (Baltimore). 2014;93(14):e72.

51. Tatsumi K, Antoniak S, Subramaniam S, Gondouin B, Neidich SD, Beck MA, et al. Anticoagulation increases alveolar hemorrhage in mice infected with influenza A. PhysiolRep. 2016;4(24).

52. Otoshi T, Kataoka Y, Nakagawa A, Otsuka K, Tomii K. Clinical Features and Outcomes of Diffuse Alveolar Hemorrhage During Antithrombotic Therapy: A Retrospective Cohort Study. Lung. 2016;194(3):475–81.

53. Kopic A, Benamara K, Schuster M, Leidenmuhler P, Bauer A, Glantschnig H, et al. Coagulation phenotype of wild-type mice on different genetic backgrounds. LabAnim. 2019;53(1):43–52.

54. Arganaraz GA, Palmeira JDF, Arganaraz ER. Phosphatidylserine inside out: a possible underlying mechanism in the inflammation and coagulation abnormalities of COVID-19. Cell CommunSignal. 2020;18(1):190.

55. Mevorach D, Mascarenhas JO, Gershov D, Elkon KB. Complement-dependent clearance of apoptotic cells by human macrophages. JExpMed. 1998;188(12):2313–20.

56. Xiang M, Jing H, Wang C, Novakovic VA, Shi J. Persistent lung injury and prothrombotic state in long COVID. FrontImmunol. 2022;13:862522.

57. Winter WE, Flax SD, Harris NS. Coagulation Testing in the Core Laboratory. LabMed. 2017;48(4):295–313.

58. Naji DH, Tan C, Han F, Zhao Y, Wang J, Wang D, et al. Significant genetic association of a functional TFPI variant with circulating fibrinogen levels and coronary artery disease. MolGenetGenomics. 2018;293(1):119–28.

59. Randi AM, Jones D, Peghaire C, Arachchillage DJ. Mechanisms regulating heterogeneity of hemostatic gene expression in endothelial cells. JThrombHaemost. 2023;21(11):3056–66.

60. Potente M, Makinen T. Vascular heterogeneity and specialization in development and disease. NatRevMolCell Biol. 2017;18(8):477–94.

61. Cleuren ACA, van der Ent MA, Jiang H, Hunker KL, Yee A, Siemieniak DR, et al. The in vivo endothelial cell translatome is highly heterogeneous across vascular beds. ProcNatlAcadSciUSA. 2019;116(47):23618–24.

62. Zhuang H, Han S, Lu L, Reeves WH. Myxomavirus serpin alters macrophage function and prevents diffuse alveolar hemorrhage in pristane-induced lupus. ClinImmunol. 2021;229:108764.

63. Jin H, Wang DY, Mei YF, Qiu WB, Zhou Y, Wang DM, et al. Mitogen-activated protein kinases pathway is involved in physiological testosterone-induced tissue factor pathway inhibitor expression in endothelial cells. Blood CoagulFibrinolysis. 2010;21(5):420–4.

64. Zhou X, Yin Z, Guo X, Hajjar DP, Han J. Inhibition of ERK1/2 and activation of liver X receptor synergistically induce macrophage ABCA1 expression and cholesterol efflux. JBiolChem. 2010;285(9):6316–26.

65. Sugiura R, Satoh R, Takasaki T. ERK: A Double-Edged Sword in Cancer. ERK-Dependent Apoptosis as a Potential Therapeutic Strategy for Cancer. Cells. 2021;10(10).

66. Nakashima K, Nishimura N, Yanagihara T, Egashira A, Ogo N, Asoh T, et al. A fatal case of diffuse alveolar hemorrhage complicated by rheumatoid arthritis. RespirMedCase Rep. 2021;32:101363.

67. Deane KD, West SG. Antiphospholipid antibodies as a cause of pulmonary capillaritis and diffuse alveolar hemorrhage: a case series and literature review. SeminArthritis Rheum. 2005;35(3):154–65.

68. Sangli SS, Ryu JH, Baqir M. Diffuse Alveolar Hemorrhage in Primary Versus Secondary Antiphospholipid Syndrome. JClinRheumatol. 2021;27(8):e297–e301.

69. Khalife WI, Kogoj P, Kar B. Sirolimus-induced alveolar hemorrhage. JHeart Lung Transplant. 2007;26(6):652–7.

70. Junpaparp P, Sharma B, Samiappan A, Rhee JH, Young KR. Everolimus-induced severe pulmonary toxicity with diffuse alveolar hemorrhage. AnnAmThorac Soc. 2013;10(6):727–9.

71. Quiroga T, Goycoolea M, Panes O, Aranda E, Martinez C, Belmont S, et al. High prevalence of bleeders of unknown cause among patients with inherited mucocutaneous bleeding. A prospective study of 280 patients and 299 controls. Haematologica. 2007;92(3):357–65.

72. Downes K, Megy K, Duarte D, Vries M, Gebhart J, Hofer S, et al. Diagnostic high-throughput sequencing of 2396 patients with bleeding, thrombotic, and platelet disorders. Blood. 2019;134(23):2082–91.

73. Mehic D, Tolios A, Hofer S, Ay C, Haslacher H, Rejto J, et al. Elevated levels of tissue factor pathway inhibitor in patients with mild to moderate bleeding tendency. Blood Adv. 2021;5(2):391–8.

74. Blann AD, Seigneur M, Steiner M, Boisseau MR, McCollum CN. Circulating endothelial cell markers in peripheral vascular disease: relationship to the location and extent of atherosclerotic disease. EurJClinInvest. 1997;27(11):916–21.

75. Yao G, Qi J, Zhang Z, Huang S, Geng L, Li W, et al. Endothelial cell injury is involved in atherosclerosis and lupus symptoms in gld.apoE(-) (/) (-) mice. IntJRheumDis. 2019;22(3):488–96.

76. Tosetto A, Castaman G, Rodeghiero F. Bleeders, bleeding rates, and bleeding score. JThrombHaemost. 2013;11 Suppl 1:142–50.

77. Bowman M, Mundell G, Grabell J, Hopman WM, Rapson D, Lillicrap D, et al. Generation and validation of the Condensed MCMDM-1VWD Bleeding Questionnaire for von Willebrand disease. JThrombHaemost. 2008;6(12):2062–6.

